# Bacteriostatic lethality emerges from turgor-induced physiological stress

**DOI:** 10.64898/2026.09.01.748669

**Authors:** Tatsuya Akiyama, Minsu Kim

**Affiliations:** Department of Physics, Emory University, Atlanta, GA, 30322. U.S.A; Graduate Division of Biological and Biomedical Sciences, Emory University, Atlanta, GA, 30322. U.S.A; Antibiotic Research Center, Emory University, Atlanta, GA, 30322. U.S.A

## Abstract

With new antibiotic discovery slowing, maximizing the efficacy of existing drugs is crucial. Bacteriostatic antibiotics are regarded as inferior to bactericidal drugs because they are thought to merely arrest growth without inflicting cellular damage and cell death. Contrary to conventional views, we show that bacteriostasis is a physiologically stressed state, which can be exploited to unmask its lethality. Through quantitative measurements and live-cell imaging, we found that translation inhibition by bacteriostatic antibiotics induces intracellular turgor pressure buildup by perturbing physiological balance in *E. coli*. This pressure manifests as osmotic swelling, leading to lysis in a subset of cells, while surviving cells depend on PBP1-mediated cell-wall repair, which alleviates mechanical strain. This protective mechanism led us to predict, and we experimentally confirmed, that inactivating PBP1 with a β-lactam amplifies the lethality of bacteriostatic drugs, challenging the general assumption of antagonism between bacteriostatic and bactericidal antibiotics. These findings show bacteriostasis as a physiologically unstable state and provide a systems-level framework for its lethality, enabling the rational design of optimized therapies.

## Introduction

Despite the success of antibiotics in modern medicine, the pipeline for development of new drugs has slowed dramatically over the past several decades. As a result, recent research has increasingly focused on improving the performance of existing antibiotics^1–3^. Although mode of action for antibiotics is well known, the cellular responses to many drugs remain incompletely understood, limiting our ability to optimize their use.

All antibiotics act through one of two fundamental modes: bactericidal agents, which kill bacteria, and bacteriostatic agents, which inhibit bacterial growth. Historically, research and clinical practice have largely favored the bactericidal drugs due to their irreversible killing^4,5^. Bactericidal antibiotics, particularly β-lactams and fluoroquinolones, are among the most widely used antibiotic classes worldwide^6,7^. Yet, their widespread use has also accelerated the emergence of resistance, which in turn undermines the killing efficacy of these drugs.

In contrast, bacteriostatic antibiotics have often been viewed as merely halting bacterial proliferation^8^. Without inflicting permanent damage like bactericidal drugs, their inhibition is transient. Once the drug is removed, bacterial growth resumes, potentially leading to relapse or re-infection. Consequently, bacteriostasis has long been regarded as a less desirable therapeutic outcome because host immunity is necessary for bacterial clearance. Although recent studies have begun to reassess their clinical potential, skepticism toward bacteriostatic agents persists, particularly in immunocompromised patients or in life-threatening infections^4^, where antibiotics are most critically needed.

This limitation has long been regarded as intrinsic, preset by the molecular mode of action of bacteriostatic drugs. For example, the most common mode of bacteriostatic action is to bind to the ribosome and inhibit translation. Because biomass synthesis drives cell growth, slowing or halting protein production directly reduces or stops growth. When translation is completely inhibited, growth ceases, resulting in bacteriostasis.

This assumption has also shaped antibiotic combination therapy. Combination therapy has reemerged as a central strategy to enhance efficacy and curb resistance^3,9^, yet bacteriostatic – bactericidal combinations are often expected to be antagonistic^10^. Bacteriostatic drugs are often excluded, under the assumption that growth arrest suppresses the cellular processes required for killing. Consistent with this view, antibiotic research and therapeutic development have historically emphasized bactericidal activity^4^.

Cell growth is an emergent outcome of many interdependent cellular processes. Perturbations that slow growth alter cell physiology in fundamental ways^11–13^. In this study, we investigated the global response of *E. coli* to bacteriostatic drugs, demonstrating that bacteriostasis is a physiologically vulnerable state, which can be exploited to unleash its lethality. Specifically, we found that translation inhibition creates a broader physiological imbalance involving metabolic imbalance, continued macromolecular accumulation, elevated turgor pressure, and cell swelling, leading to partial lysis in *E. coli*. Using genetic knock-out and downregulation, we reveal that this physiological vulnerability is typically masked by pressure relief by mechanosensitive channels, and particularly by cell wall repair system by PBP1. Consequently, inhibiting PBP1 with a β-lactam unmasked this vulnerability, enhancing the killing by bacteriostatic drugs, contrary to the general assumption of bacteriostatic – bactericidal antagonism. These results demonstrate a mechanism for “bacteriostatic lethality,” offering a novel bacteriostatic-centric perspective for improving antibiotic therapy.

## Results

### Bacteriostatic drugs induce complex physiological perturbations

Protein synthesis is a primary driver of cell growth and therefore the most common target of bacteriostatic drugs^8^. Yet, translation is also among the most energetically demanding cellular processes^14^, tightly coupled to metabolism^12^. To characterize the effect of translation inhibition on cell physiology, we monitored *E. coli* growth under chloramphenicol treatment. As chloramphenicol concentration increases, culture growth ultimately stops (Fig. 1A). The lowest concentration required to stop growth was defined as the minimum inhibitory concentration (MIC = 40 µg/mL).

**Figure 1.**
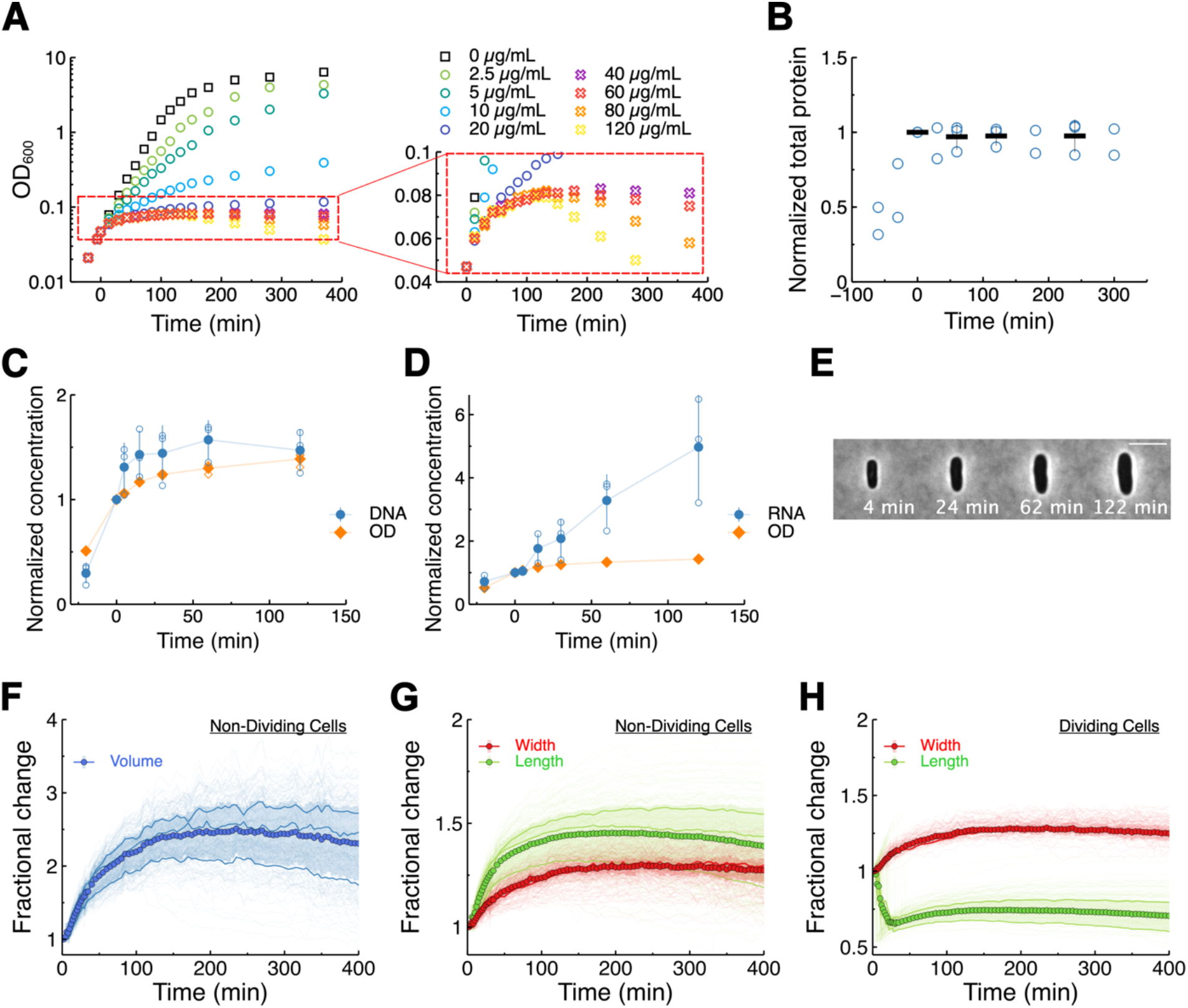
Cellular growth persists after acute translation inhibition. (A) Optical density (OD) of chloramphenicol-treated culture increases even at high concentrations. (B) Total protein stops increasing immediately after chloramphenicol treatment. Black bars represent the average of three biological replicates. (C) DNA and (D) RNA concentrations continue to increase during chloramphenicol treatment. Data represent three biological replicates. (E) Microscope images show cellular expansion of chloramphenicol-treated cells. Chloramphenicol-treated cells increase in (F) volume and (G) length and mean width. (H) Cells that complete division after treatment also increase in mean width. Thin lines represent individual cells (233 dividing and 328 non-dividing cells), bold lines represent the mean of each biological replicate, and markers represent the mean of the three biological replicates.

Unexpectedly, at the MIC, optical density (OD) nearly doubled before plateauing, indicating substantial post-treatment growth. Even at concentrations exceeding the MIC, this residual increase persisted (Fig. 1A). To test whether this residual growth was caused by delayed translation inhibition after antibiotic addition, we measured total protein using the Biuret method. The results indicate that inhibition of protein synthesis above the MIC is immediate (Fig. 1B).

We next asked whether translation inhibition also stopped the accumulation of other major macromolecules. Despite the rapid arrest of protein synthesis, total DNA and RNA continued to increase during the post-treatment period (Fig. 1CD). Thus, translation arrest does not immediately stop all macromolecular accumulation, as non-protein biomass components continue to accumulate after protein synthesis is arrested.

The continued OD increase could simply reflect residual normal growth that gradually slows after treatment. Under normal growth conditions, cell length increases while cell width remains constant^15^. However, single-cell time-lapse imaging revealed that the post-treatment growth did not follow this normal growth pattern. Instead, elongation slowed while cell width increased, causing cell volume to expand for extended periods after treatment (Fig. 1E-G, Movie 1). Cells already undergoing division often completed the division (Fig. 1H). Similar post-treatment expansion was observed for other translation inhibitors, including tetracycline and erythromycin (Extended Data Fig. 1). Together, these results demonstrate that translation inhibition does not immediately arrest macromolecular accumulation or cell growth but triggers a transient phase of abnormal morphological change and continued volume expansion.

### An increase in turgor pressure results in osmotic swelling

The observation that cells continue to expand (Fig. 1A, E-G) despite immediate translation arrest (Fig. 1B) is striking, as translation is normally the principal driver of biomass accumulation and therefore of growth. This decoupling between protein synthesis and volume increase suggests a mechanism of cell expansion distinct from typical translation-driven growth. Because turgor pressure mechanically loads the cell envelope and can drive cell expansion, we asked whether translation inhibition perturbs intracellular turgor.

Turgor pressure arises from the osmotic difference between the internal and external environment^16^. Typically, when the external osmolarity exceeds the internal osmolarity in osmotic ramp-up, water efflux causes the cytoplasm to shrink and detach from the rigid cell wall, a process known as plasmolysis. The external osmolarity increase required to trigger plasmolysis provides a quantitative measure for the turgor pressure^17–19^.

In our experiments, we trapped cells in a microfluidic chamber, increased the external osmolarity by adding higher concentrations of NaCl, and monitored plasmolysis using a microscope (Fig. 2A). Untreated cells plasmolyzed at or below an additional 200 mM NaCl, whereas chloramphenicol-treated cells required 300 mM or higher NaCl to plasmolyze (Fig. 2B), indicating elevated turgor pressure in translation-inhibited cells.

**Figure 2.**
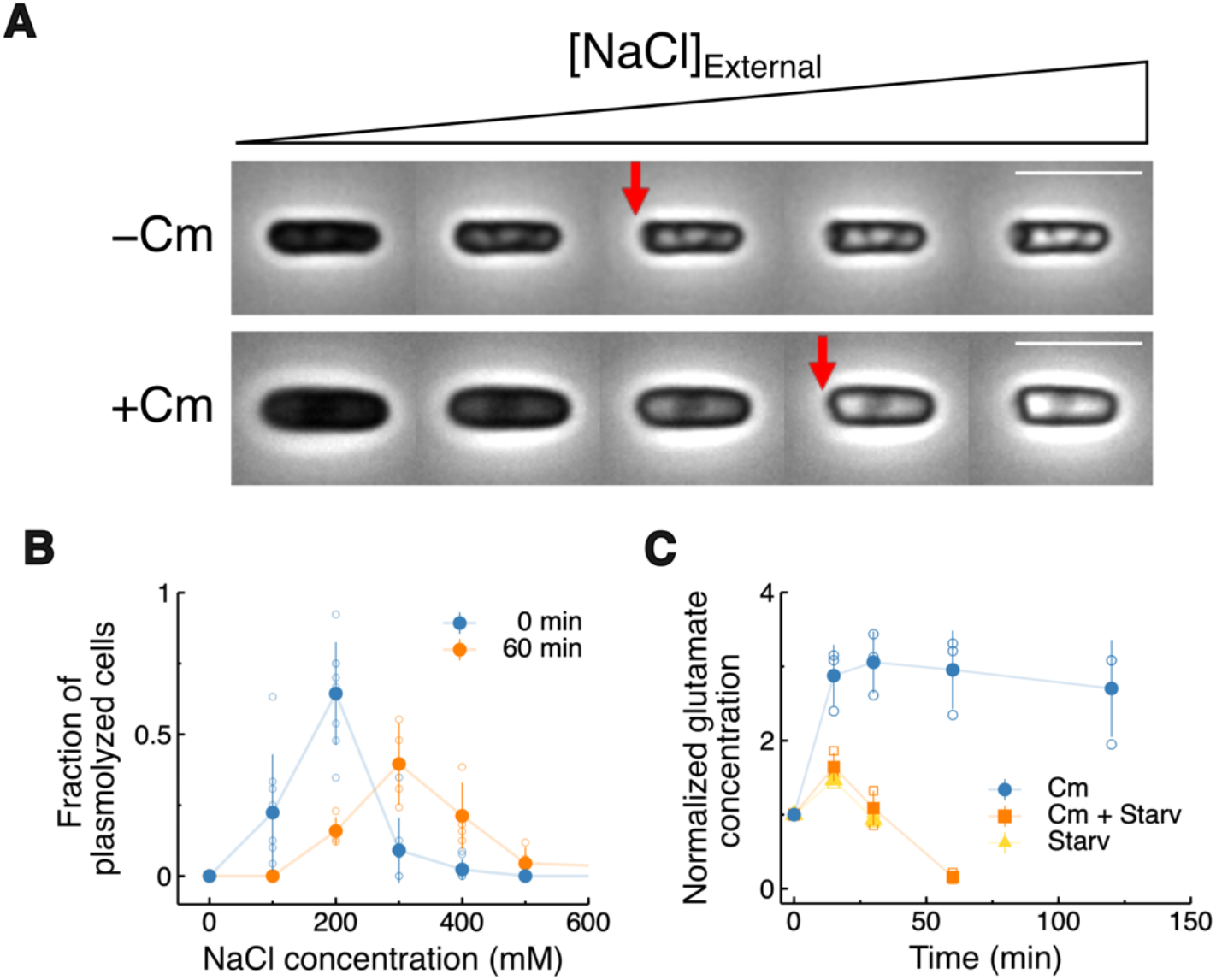
Continued metabolism elevates cell-associated glutamate and turgor pressure after translation arrest. (A) Microscope images showing plasmolysis induced by increasing external NaCl concentrations. Red arrow indicates plasmolysis. (B) Chloramphenicol-treated cells require higher external osmolarity to induce plasmolysis. Data represent four to eight biological replicates at each NaCl concentration. (C) Cell-associated glutamate concentration increases during chloramphenicol treatment when nutrients are available. Data represent three biological replicates.

We further tested this by holding the external osmolarity constant at 200 mM additional NaCl on agarose pads and quantifying the fraction of plasmolyzed cells. The majority of untreated cells plasmolyzed under these conditions, whereas chloramphenicol-treated cells exhibited only a small fraction of plasmolyzed cells (Extended Data Fig. 2). Together, these results demonstrate that translation inhibition elevates intracellular turgor pressure.

### Nutrient utilization contributes to intracellular osmolyte accumulation

We next investigated how translation inhibition increases turgor pressure. Intracellular metabolites are the primary determinants of cytoplasmic osmolarity and hence of turgor pressure^16^. In *E. coli*, glutamate is one of the most abundant organic anions, serving both as a major osmolyte itself and as the major counterion to potassium, the dominant cationic osmolyte^20–22^. When we measured the intracellular glutamate concentration, it increased by 3-fold following translation inhibition (Fig. 2C).

Because no known regulatory pathway directly links translation inhibition to glutamate synthesis, we reasoned that this increase likely reflects passive accumulation. Continued nutrient utilization after translation inhibition could contribute to metabolite accumulation because metabolites continued to be produced while their incorporation into proteins is blocked. Such imbalance should elevate a broad range of metabolites. Indeed, previous metabolomic studies report chloramphenicol-induced increases in proline, glycine, and other compatible osmolytes^23,24^. In the next experiment, we deprived translation-inhibited cells of nutrients to suppress nutrient utilization. Intracellular glutamate levels barely increased (Fig. 2C), and cell width no longer increased (Extended Data Fig. 3). Together, these findings suggest that sustained nutrient utilization contributes to intracellular osmolyte accumulation, elevated turgor pressure, and width expansion after translation inhibition. This pressure-driven expansion is mechanistically distinct from balanced growth and is consistent with osmotic swelling.

### Osmotic swelling promotes cell lysis

In our study, chloramphenicol treatment caused a 2.5-fold increase in cell volume (Fig 1F), far exceeding the approximately 15% expansion reported after hypotonic shock^25^ and indicating that passive turgor-driven stretching of the preexisting envelope alone cannot account for the observed volume increase. To evaluate its effect on the envelope damage, we used the membrane-impermeant dye SYTOX Green, which enters cells only when the envelope is compromised. Following chloramphenicol treatment, SYTOX Green fluorescence increased in a subset of cells, indicating loss of membrane integrity (Extended Data Fig. 4). In these cells, fluorescence often increased sharply and was accompanied by cell shrinkage, consistent with rupture and lysis (Extended Data Fig. 4).

The frequency of cell death increased with higher chloramphenicol concentrations. At the MIC (40 µg/mL), approximately 1% of cells lost viability after 5 hours (Fig. 3C). A modest increase in concentration caused a marked rise in killing. Increasing the concentration from 40 to 60 µg/mL increased the fraction of dead wild type cells (Fig. 3C; blue symbol; *p* < 0.05, two-way ANOVA followed by Fisher’s LSD). These findings are consistent with a decrease in OD observed after translation inhibition, which became steeper at higher chloramphenicol concentrations (Fig. 1A). These results show that the turgor pressure build-up under translation inhibition compromises envelope integrity and causes lysis in a subset of cells.

**Figure 3.**
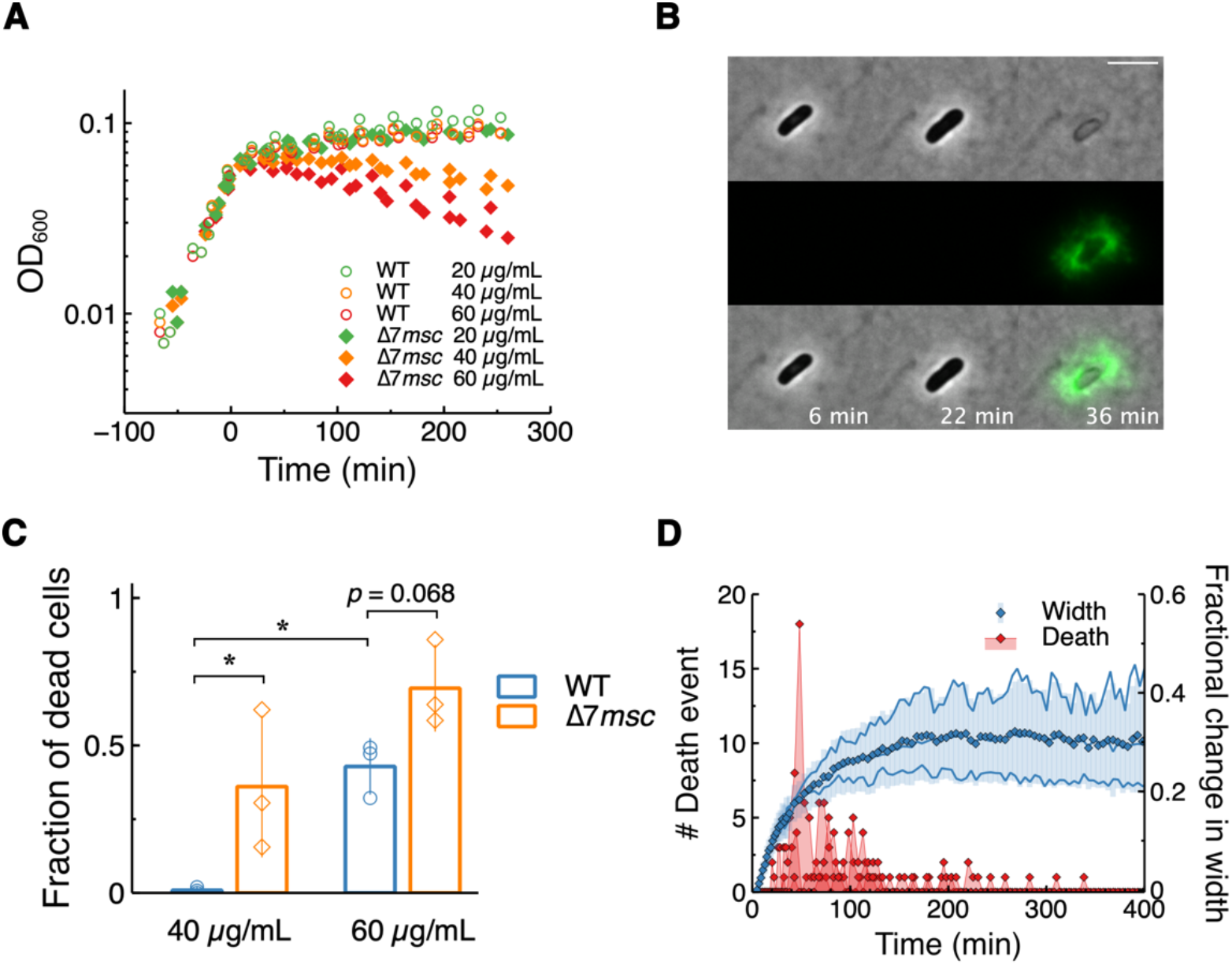
Elevated turgor pressure during chloramphenicol-induced cellular expansion promotes cell lysis. (A) Loss of mechanosensitive channels (Δ7*msc*) sensitizes cells to chloramphenicol as shown by OD decrease. Data represent three biological replicates. (B) Microscopy shows chloramphenicol-induced cell lysis detected using the membrane-impermeable dye SYTOX Green. (C) Loss of mechanosensitive channels increases susceptibility to chloramphenicol-induced cell lysis. Data represent three biological replicates with 325 to 630 cells analyzed per condition. (D) Lysis occurs predominantly during the period of rapid cell-width expansion during treatment with 40 µg/mL chloramphenicol in the absence of mechanosensitive channels. Thin lines represent individual cells (437 cells total), bold lines represent the mean of each biological replicate, and markers represent the mean of the three biological replicates.

To further evaluate the role of turgor in cell death, we tested whether a turgor-sensitive mutant strain is sensitive to chloramphenicol treatment. Mechanosensitive channels (MSCs) are safety valves that open when membrane tension reaches a critical threshold, allowing solute efflux to relieve excess turgor^26^. Mutants lacking MSCs are predicted to experience higher internal pressure upon osmotic downshift, as inferred from their excessive swelling and lysis^25^. Our results showed that even if the external osmolarity does not change, under chloramphenicol treatment, cells experience higher internal pressure. This is expected to be more severe for a strain with seven *msc* genes deleted (Δ7*msc*), exacerbating the lysis.

The Δ7*msc* mutant displayed a residual increase in OD following chloramphenicol treatment, similar to the wild type, consistent with osmotic swelling. However, this was subsequently followed by a steeper decline, indicating extensive cell lysis (Fig. 3A). This was confirmed by single-cell microscopy, which revealed a higher frequency of cell death in Δ7*msc* cells compared with the wild type (Fig. 3BC; *p* < 0.05 and *p* = 0.068 for 40 and 60 µg/mL chloramphenicol, respectively; two-way ANOVA followed by Fisher’s LSD). Lysis occurred most frequently during the initial swelling period (Fig. 3BD), underscoring the detrimental effect of the pressure buildup. Under nutrient deprivation which prevents the pressure increase (Extended Data Fig. 3), Δ7*msc* mutant showed greatly reduced lysis (Extended Data Fig. 5). These findings further support our conclusion that an increased turgor pressure under translation inhibition is mechanically harmful to the cells.

### PBP1-dependent cell-wall synthesis protects cells during swelling

Despite substantial width expansion and an approximately 2.5-fold increase in cell volume, a large fraction of cells did not lyse. How do cells survive this turgor-driven swelling? Because expansion under turgor requires cell envelope remodeling and synthesis^18,27^, we asked whether cell wall synthesis continued after translation inhibition. Although reduced, cell wall synthesis continued in translation-inhibited cells (Fig. 4A), indicating that cellular expansion is accompanied by ongoing cell wall synthesis.

**Figure 4.**
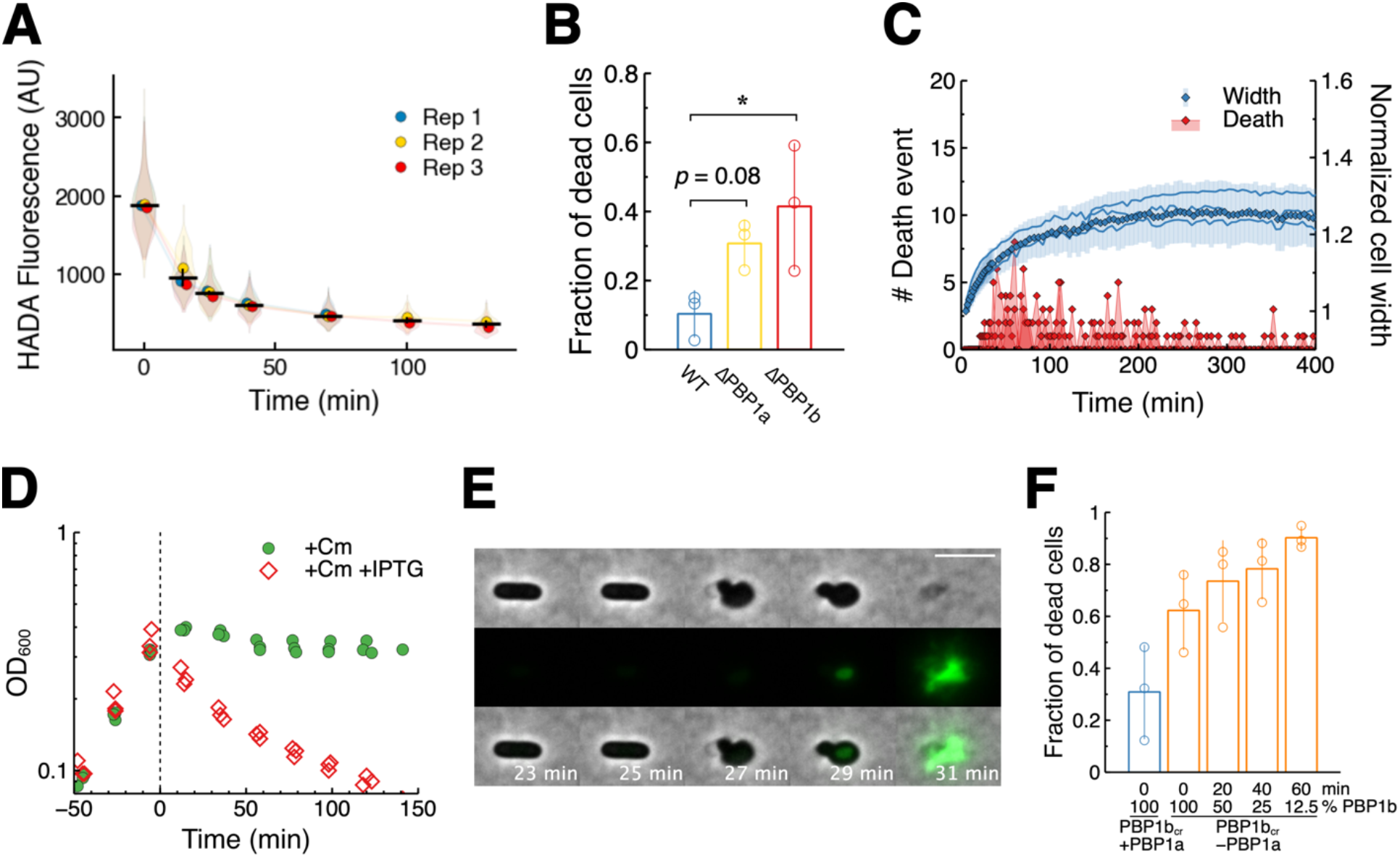
Residual cell wall synthesis by PBP1 during chloramphenicol-induced cellular expansion protects cells from lysis. (A) Cell wall synthesis continues at a reduced rate during chloramphenicol treatment as shown by HADA incorporation. Data represent two to three biological replicates with 326 to 917 cells analyzed per condition. (B) Deletion of either PBP1a or PBP1b sensitizes cells to chloramphenicol, as measured by microscopy. Data represent three biological replicates with 515 to 598 cells analyzed per condition. (C) Chloramphenicol-induced lysis of ΔPBP1b cells occurs during cellular expansion. Thin lines represent individual cells (517 cells total), bold lines represent the mean of each biological replicate, and markers represent the mean of the three biological replicates. (D) PBP1b was depleted by IPTG-inducible CRISPRi in a PBP1a deletion background. PBP1 depletion sensitizes cells to chloramphenicol as shown by the decrease in OD. (E) Microscopy images showing abrupt cell lysis of PBP1-depleted cells during chloramphenicol treatment. (F) Chloramphenicol-induced cell lysis increases as PBP1 is depleted. Data represent three biological replicates with 928 to 1111 cells analyzed per condition.

In *E. coli* and many other bacteria, three major transpeptidases contribute to peptidoglycan synthesis: penicillin-binding proteins (PBPs) 1, 2, and 3^28^. PBP2 and PBP3 are the principal enzymes associated with elongation and division, respectively, during active growth, whereas recent evidence suggests that PBP1 primarily functions in cell wall repair^29–31^. Another interesting distinction is that PBP2- and PBP3-mediated synthesis is highly directional, maintaining cell shape and constant width^32,33^, whereas PBP1 activity appears less constrained. Indeed, a previous study showed that increased PBP1 activity leads to cell widening in *Bacillus subtilis*^34^. In our experiments, translation-inhibited cells also exhibited a marked increase in width during residual growth and ongoing cell wall synthesis (Fig. 1GH, Fig. 4A). Based on these considerations, we became curious about a potential role of PBP1 in translation-inhibited cells.

*E. coli* encodes two partially complementary PBP1 enzymes, PBP1a and PBP1b^31^. Single knockouts of either gene are viable under normal conditions. When treated with chloramphenicol, both mutants showed a pronounced increase in cell lysis compared with the wild type (Fig. 4B), indicating that PBP1 activity is critical for tolerance to chloramphenicol-induced cell lysis. Notably, we observed pronounced cell death (Fig. 4C; diamonds, left axis) during the residual width increase (lines, right axis), suggesting that PBP1 plays an important role in protecting cells against turgor-induced rupture during osmotic swelling.

To further evaluate this role of PBP1, we wished to knock out both PBP1a and 1b, but the double knockout is lethal. We therefore used a CRISPR interference (CRISPRi) system to repress PBP1b expression on demand in the ΔPBP1a background. Upon induction of PBP1b repression, preexisting PBP1b supported normal exponential growth for approximately four doublings, after which growth ceased (Extended Data Fig. 6). In a parallel culture, we added chloramphenicol during this exponential growth phase: about 2.5 doublings after PBP1 repression, where we estimate PBP1b levels had decreased to ∼17% of the original amount (1/2^2.5^). Chloramphenicol caused an immediate and steep decline in OD (Fig. 4D), in contrast to the control (no PBP1b repression), which exhibited only a minor decline.

This hypersensitivity of PBP1-depleted cells to lysis was also evident under the microscope. Upon chloramphenicol treatment, some PBP1b-depleted cells abruptly transitioned from a rod shape to a rounded morphology before lysis (Fig. 4E), indicating that the cells could no longer maintain rod shape against the rising turgor pressure. This deformation was immediately followed by a sudden pop, corresponding to rapid lysis (Fig. 4E). To further evaluate the effect of PBP1, we added chloramphenicol at progressively later times after initiating PBP1b repression, corresponding to lower estimated PBP1b level ranging from 12.5 – 100%. We found that the frequency of lysis increased as PBP1b levels decreased (Fig. 4F). Together, these results demonstrate a critical role of PBP1 in cell survival during translation inhibition.

### PBP1-targeting β-lactam potentiates translation-inhibiting antibiotics

The critical role of PBP1 in withstanding turgor-induced mechanical stress predicts that inactivating PBP1 with β-lactams should potentiate chloramphenicol-induced lysis. Importantly, this prediction challenges the general expectation of static-cidal antagonism. To test this prediction, we added the MIC of PBP1-targeting β-lactam cefsulodin (48 µg/ml) to chloramphenicol-treated cultures. Consistent with our hypothesis, OD declined rapidly after cefsulodin addition (Fig. 5A). We confirmed this decline is due to extensive cell lysis using single-cell microscopy (Fig. 5B).

**Figure 5.**
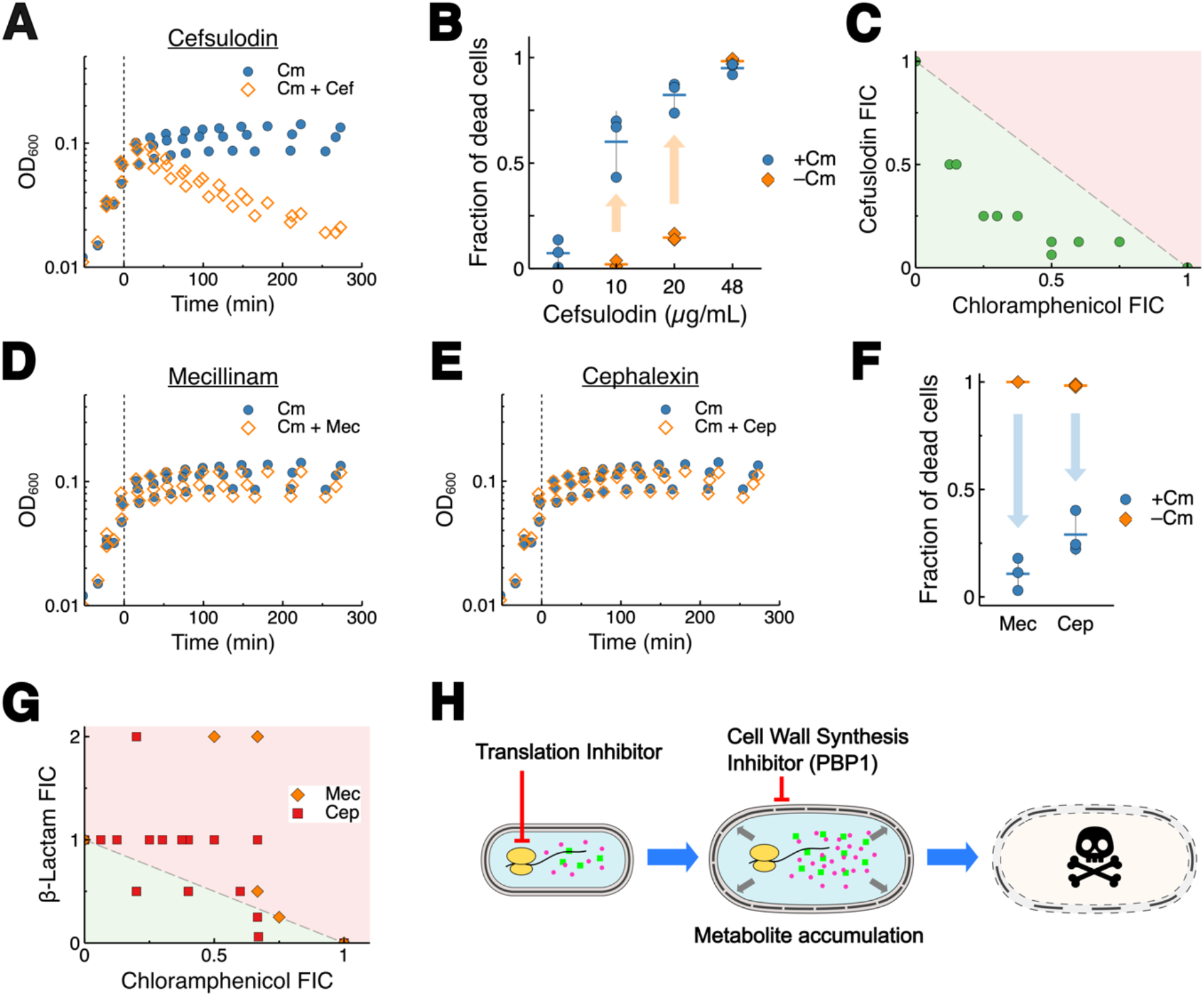
PBP1-targeting β-lactam enhances chloramphenicol-induced bactericidal activity. (A) Chloramphenicol-treated cells are sensitive to the PBP1-targeting β-lactam cefsulodin as shown by decreased OD. (B) A nonlethal dose of cefsulodin potentiates chloramphenicol-induced cell lysis measured by microscopy. Data represent three biological replicates with 566 to 994 cells analyzed per condition. (C) A checkerboard assay shows a positive interaction between cefsulodin and chloramphenicol at sub-inhibitory concentrations. Unlike cefsulodin, (D) PBP2-targeting mecillinam and (E) PBP3-targeting cephalexin did not kill chloramphenicol-treated cells. (F) Single-cell measurements and (G) checkerboard assay show antagonistic interactions between chloramphenicol and mecillinam/cephalexin. For (F), data represent three biological replicates with 355 to 1048 cells analyzed per condition. (H) Proposed model in which continued cellular expansion after translation inhibition increases dependence on residual PBP1-mediated cell wall synthesis. PBP1 inhibition promotes turgor-driven cell lysis.

Our PBP1 depletion experiments above suggested that even partial inhibition of PBP1 was sufficient to trigger cell death in translation-inhibited cells (Fig. 4F). We therefore examined sublethal concentrations of cefsulodin, which alone had no effect on cell lysis in normally growing cells (Fig. 5B, Extended Data Fig. 7A). When the same sublethal concentrations were applied to translation-inhibited cells, pronounced lysis occurred (Fig. 5B, Extended Data Fig. 7B), revealing a strong positive interaction between translation-inhibiting and PBP1-targeting drugs.

We further quantified drug interactions using a checkerboard assay. In this assay, cells were grown in media containing varying concentrations of two drugs arranged in a 96-well plate. From the resulting growth profiles, we determined the MICs for each drug alone and in combination and calculated their fractional inhibitory concentrations (FICs). The sum of the two FICs, FIC index (FICi), provides a quantitative measure of drug interactions^35^. Specifically, an FICi of 1 indicates no interaction (additivity; dashed line in Fig. 5C), an FICi greater than 1 indicates negative interaction (red), and an FICi less than 1 indicates positive interaction (green). For the chloramphenicol – cefsulodin combination, the FICi for all combinations at sublethal concentrations were below 1, showing positive interactions (Fig 5C). Similar positive interactions with cefsulodin were also observed for other translation inhibiting bacteriostatic drugs (tetracycline and erythromycin) (Extended Data Fig. 8).

We next evaluated the effects of the other two PBPs. Mecillinam and cephalexin preferentially inactivate PBP2 and PBP3, respectively^36^. In contrast to cefsulodin, adding either of these drugs at their MICs to translation-inhibited cultures failed to produce any decline in OD (Fig. 5DE). Single-cell microscopy confirmed that, when used alone, these β-lactams lysed the majority of growing cells, but when combined with chloramphenicol, the frequency of lysis dropped dramatically (Fig. 5F), indicating antagonism. This antagonism was further supported by checkerboard assays, in which FICi values remained above 1 (Fig. 5G). Thus, the positive interaction observed with chloramphenicol is specific to PBP1-targeting β-lactams.

## Discussion

The conventional view holds that bacteriostatic antibiotics merely halt growth without causing cellular damage. Our findings challenge this view. We show that bacteriostasis creates a broader physiological imbalance involving metabolic imbalance and continued macromolecular accumulation, leading to elevated turgor pressure. This elevated turgor pressure drives abnormal width expansion and cell swelling that are mechanically distinct from normal cell growth, a process we term osmotic swelling (Fig. 5H). This mechanical vulnerability is evident in the death of a subset of cells, whose frequency increases with higher drug concentrations. The vulnerability becomes fully exposed when the cell-wall repair enzyme PBP1 is inactivated, leading to complete loss of structural integrity and subsequent lysis. This finding establishes *bacteriostatic lethality* as a systems-level emergent property arising from physiological imbalance that ultimately leads to structural failure and death.

Dating back to the 1950s and continuing to recent years, there have been reports translation-inhibiting bacteriostatic drugs can lead to loss of cell viability^37–39^. Interestingly, several of these studies also suggested cell-envelope damage^38,40^. Yet, their underlying mechanisms remained unresolved. Our finding of *bacteriostatic lethality* fills this gap by providing a systems-level explanation for how translation inhibition can convert growth arrest into cell death via the envelope damage.

More broadly, our findings highlight that antibiotic action cannot be understood solely in terms of the primary molecular target. The activity of translation-inhibiting antibiotics is also strongly dependent on bacterial growth state, with susceptibility varying with growth rate^11^ and chloramphenicol killing occurring preferentially in rapidly growing cells^38^. Several antibiotics exhibit dual or shifting modes of action, acting as bacteriostatic under some conditions and bactericidal under others, even though their molecular targets remain unchanged. For example, nalidixic acid can produce either bacteriostatic or bactericidal effects depending on the cellular processes and respiratory state^41–43^. Trimethoprim and sulfonamides, each bacteriostatic on their own, can become bactericidal when combined^44^. Recent work has also suggested that some bacteriostatic antibiotics, while nonlethal in standard laboratory conditions, might become bactericidal inside host environments where stress responses or metabolic states differ. For example, azithromycin, a primarily bacteriostatic translation inhibitor, can exhibit bactericidal activity under physiological conditions that better mimic the host environment^45^. It remains unclear why the same molecular inhibition can lead to such divergent outcomes. Our systems-level framework for antibiotic action highlights that antibiotic activity is not a fixed property of its molecular target but a context-dependent consequence of the cell’s multidimensional physiological state.

This framework can help maximize the efficacy of existing antibiotics. Our prediction of combination therapy was bacteriostatic-centric, predicting how the lethality of translation-inhibiting bacteriostatic drugs is enhanced by the β-lactam targeting PBP1. This prediction runs counter to the prevailing bactericidal-centric view that bactericidal activities are inhibited by static drugs. The experimental verification of our prediction demonstrates how our new perspective can reveal unexpected opportunities to enhance antibiotic potency.

Conflicting reports of synergy and antagonism among antibiotics within the same class, particularly β-lactams, have persisted for decades. Drugs sharing similar chemical scaffolds or targets often produce divergent outcomes, with some combinations enhancing killing and others suppressing it, complicating our predictive understanding of combination therapy^46,47^. Our findings provide insight into this puzzling phenomenon. PBP2 and PBP3 are responsible for cell-wall synthesis during elongation and division, processes that are inactivated when translation is inhibited. Consistent with the prevailing hypothesis, we found that translation inhibition abolishes the killing activity of β-lactams targeting PBP2 or PBP3, leading to antagonism. By contrast, PBP1 functions primarily in cell wall repair, which explains why PBP1-targeting β-lactam potentiates the effects of translation inhibition, as discussed above. Importantly, all of these PBP-targeting drugs are categorized as β-lactams. These comparisons provide a physiological explanation for how antibiotics with shared chemical scaffolds can yield opposite interaction outcomes, depending on how their specific targets engage the stressed cellular state.

These findings also have implications for drug discovery. Target-based antibiotic discovery strategies prioritize essential genes and molecular functions required for bacterial survival^1,48^. However, our results show that lethality often emerges not from the primary target itself, but from the broader physiological network. Incorporating such systems-level interactions into antibiotic discovery could reveal new vulnerabilities and expand the landscape of viable targets beyond the traditional essential-gene paradigm.

## Materials and Methods

### Bacterial strains and growth conditions

*E. coli* strains (Supplemental Table 1) were grown in LB broth (Miller) supplemented with 10 mM glucose and 1 mM MgSO_4_ (supplemented LB). Briefly, a single colony was inoculated into media in borosilicate glass culture tubes and incubated at 37°C with shaking (250 rpm) in a water bath shaker overnight. Next morning, optical density (OD_600_) was measured using a Genesys20 spectrophotometer (Thermo Fisher) with a standard cuvette (16.100-Q-10/Z8.5, Starna Cells Inc). The culture was diluted with fresh media to OD_600_ of ∼0.001 and incubated in a water bath shaker at 37°C with shaking. When cultures reached OD_600_ 0.040 – 0.060, antibiotics or IPTG were added. Strains carrying *mrcA* (PBP1a) or *mrcB* (PBP1b) deletions were generated using P1 transduction^1^.

### Time lapse imaging of bacterial growth

Time lapse microscopy was performed as previously described with modifications^2^. When the culture reached OD_600_ > 0.2, cells were placed onto a coverslip and covered with a 1 % agarose pad prepared in supplemented LB containing 1 µM SYTOX Green (Invitrogen™) and antibiotics. Another coverslip was placed on top of the agarose pad to reduce drying. Cells were imaged every 2 min for the first 30 min and then every 5 min for the rest of experiments using an inverted fluorescence microscope (Olympus IX83) with an oil immersion phase-contrast 60x objective seated inside an incubator chamber (InVivo Scientific) pre-warmed to 37°C. Images were captured using a Neo 5.5 sCMOS camera (Andor). The microscope was controlled with MetaMorph software (Molecular Devices). For endpoint measurements of live fractions, cells were incubated at 37°C and imaged at the end of incubation period. Cells that accumulated SYTOX Green fluorescence or lysed, producing fluorescent smear, were considered dead.

### Image analysis

Masks of cell shape were generated from phase contrast image using omnipose^3^. Mask was used to generate contours of cells and morphological features were measured using a MicrobeJ plugin for ImageJ^4,5^. Background of fluorescence images were subtracted using a rolling ball function and mean intensity per cell was measured by MicrobeJ. Fluorescence measurement was used to determine cell lysis. Cell size was normalized to initial time point, which is less than 8 min of exposure. Cell volume was calculated from length (l) and mean width (w) using a formula described previously^6^:

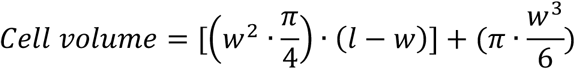

### Total protein measurement

The Biuret method described by Hui et al.^7^ was used to quantify total protein in cell cultures during the exponential phase. Briefly, 1.8mL of cell culture was collected by centrifugation at 10,000 x g for 2 min and resuspended in 0.2mL of ultra-pure water and frozen on dry ice until the assay. After thawing in water bath at room temperature, 0.1mL of 3M NaOH was added to the cells and the samples were incubated at 100°C for 5 min to hydrolyze proteins, followed by cooling at room temperature for 5 min. To carry out the Biuret reaction, 0.1mL of 1.6% CuSO_4_ was added to the samples and mixed thoroughly for 5 min at room temperature. After centrifugation, the absorbance at 555 nm was measured using a Genesys20 spectrophotometer (Thermo Fisher). A standard curve was generated by applying the same Biuret reaction to a series of BSA standards, and the protein amounts in the samples were determined by reference to the standard curve.

### DNA measurement

A 200 µL aliquot of culture was spun down at 10,000 x g for 1 min. After removing 190 µL of the supernatant, the cell pellet was lysed with 100 µL of BugBuster^®^ Protein Extraction Reagent (MilliporeSigma). DNA concentration in the crude lysate was measured using the Qubit™ dsDNA HS Assay Kit (Invitrogen™) and a BioTek Synergy H1 Multimode Reader (Agilent).

### RNA measurement

A 200 µL aliquot of culture was centrifuged and total RNA was extracted using TRIzol™ Reagent (Invitrogen™) according to the manufacturer’s instructions. Following phase separation, 300 µL of the aqueous phase was collected for RNA precipitation. RNA pellets were dissolved in 30 µL of DEPC-treated water. RNA concentration was measured using the AccuBlue^®^ Broad Range RNA Quantification Kit (Biotium) and a BioTek Synergy H1 Multimode Reader (Agilent).

### Glutamate measurement

Two tubes with 1mL of culture in supplemented LB or in 1.16% NaCl were spun down at 10,000 x g for 1 min. 950 µL of supernatant was removed. Cell pellet was washed with 950 µL of 1.16% NaCl once. After removing 950 µL of supernatant, cell pellet was resuspended in leftover solution (∼50 µL). Cell suspension was mixed with 250 µL of −80°C methanol and vortexed for 5 sec. Samples were stored at −80°C. Cells were lysed by three freeze – thaw cycles in liquid nitrogen and room temperature with 5 sec vortex every cycle. Lysate was centrifuged at 15,000 x g for 10 min. Supernatant was transferred to a new tube and dried in speed vac. Dried pellets from two tubes were resuspended in total of 50 µL water.

Glutamate concentration was determined using a Glutamate Assay Kit (MilliporeSigma) following supplier’s protocol. Absorbance at 565nm was measured using a 96 well plate (Corning) and a BioTek Synergy H1 Multimode Reader (Agilent).

### Quantification of plasmolyzed cells

Cells were plasmolyzed on custom fluidic chips constructed from a double-sided adhesive sheet and two coverslips. A larger coverslip was cleaned with 3 M HCl overnight and rinsed with water. After air drying, a coverslip was coated with 0.01% poly-L-lysine (MilliporeSigma). A double-sided adhesive sheet with channels cut out was sandwiched between a coated coverslip and a non-coated smaller coverslip. Cells were loaded onto a chip and incubated at 37°C for 15 min to allow them to adhere to the surface. For chloramphenicol-treated cells, this time is also counted as treatment time. Flow was generated by placing kimwipe or paper towel on outlet side of the chip. The chip was flushed with supplemented LB containing chloramphenicol to wash off unattached cells. Cells were imaged while flushing the chip with supplemented LB containing increasing concentrations of NaCl. Cells were also plasmolyzed on 1% agarose pads prepared in supplemented LB containing 200 mM NaCl.

### HADA labeling of cell wall

A 500 µL aliquot of culture was transferred to a 2 mL tube and incubated with 100 nM HADA (Bio-Techne/Tocris) at 37°C with shaking at 200 rpm for 20 min. Cells were fixed by directly adding 1mL of ice-cold 100% ethanol and incubating at 4°C for 20 min. Fixed cells were washed three times with 1x PBS. Cells were imaged between a 1% agarose pad prepared in PBS and a coverslip using a DAPI filter set. Average maximum HADA fluorescence was measured for individual cells by drawing multiple perpendicular line profiles from the cell centerline using ImageJ and Python. Population averages from biological replicates and single-cell fluorescence distributions are reported.

### Checkerboard assay and growth measurements in 96-well plates

Drug-drug interaction was determined by a checkerboard assay. Checkerboard plates were prepared by placing 100µL of supplemented LB containing 2x concentrated drug 1 and 50 µL of supplemented LB containing 4x concentrated drug 2 onto 96 well plates (Corning). When OD_600_ of bulk culture reached >0.3, cell density was adjusted to OD_600_ 0.001 in supplemented LB and 50 µL of cells was plated. For endpoint measurements, plates were incubated at 37°C for 16 – 24 hours. Plates were briefly shaken and OD_600_ was measured using a BioTek Synergy H1 Multimode Reader (Agilent). For time-course growth measurements, plates were continuously shaken and OD_600_ was measured every 10 min at 37°C.

The fractional inhibitory concentration (FIC) index was determined by the calculation described by Hallander et al^8^.

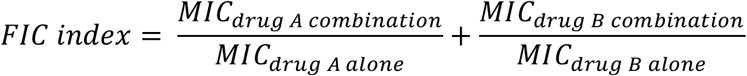

### Statistical Analysis

Welch’s two-tailed t-test was used for pairwise comparisons. For datasets with more than two groups, one-way or two-way ANOVA followed by Fisher’s LSD test was used for planned comparisons. All statistical analyses were performed using Python and Excel.

## Acknowledgements

We thank Teuta Pilizota for providing the Δ7*msc* strain and Kerwyn C. Huang for providing the CRISPRi strain. This work was funded by NIH (1U19AI158080), Human Frontier Science Program (RGY0072/2015), MP3 Initiative (00097584) and Keck foundation (0000070212).

## Author Contributions

TA and MK conceived the study. TA designed, carried out the experiments, analyzed and interpreted the data. MK secured funding and provided resources. TA and MK wrote the manuscript. All authors read and approved the manuscript.

