## Supplemental information for "Bacteriostatic lethality emerges from turgor-induced physiological stress"

Supplemental Table 1. Bacterial strains used in this study.

| Strain | Genotype | Relevant features | Source/Reference |
|---|---|---|---|
| NMK1 | | NCM3722 | |
| GMK265 | | BW25113 | |
| GMK434 | *ΔmscS::frt*, *ΔmscL::frt*, *ΔkefA::frt*, *ΔybdG::frt*, *ΔyjeP::frt*, *ΔybiO::frt*, *ΔynaI::frt* | Deletion of 7 mechanosensitive channels (*Δ7msc*) on BW25113 | Gift from Teuta Pilizota |
| JW3359 | *ΔmrcA::km* | ΔPBP1a on BW25113 | Keio collection |
| JW0145 | *ΔmrcB::km* | ΔPBP1b on BW25113 | Keio collection |
| NMK424 | *ΔmrcA::km* | ΔPBP1a on NCM3722 | This study |
| NMK422 | *ΔmrcB::km* | ΔPBP1b on NCM3722 | This study |
| 5A-WellG12-mrcB | $PBP1b_{cr}$ | CRISPRi targeting *mrcB* on BW25113 | Silvis et al. 2021 |
| GMK311 | *ΔmrcA::km*, $PBP1b_{cr}$ | ΔPBP1a, CRISPRi targeting *mrcB* on BW25113 | This study |

Silvis, M. R. *et al.* Morphological and Transcriptional Responses to CRISPRi Knockdown of Essential Genes in Escherichia coli. *mBio* **12**, e0256121 (2021).

Supplemental video 1. Morphological change of *E. coli* cell during chloramphenicol treatment.

Extended data figures

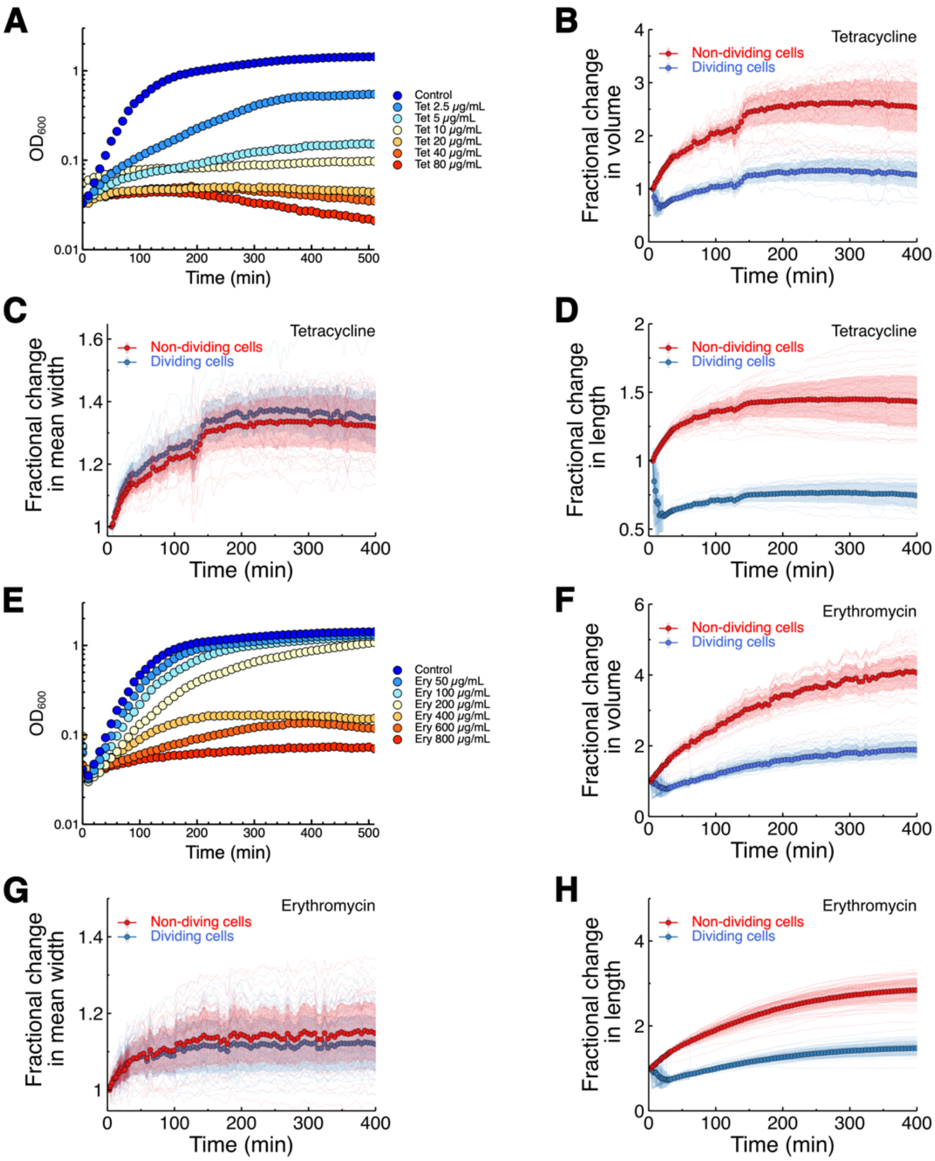


Extended Data Fig 1. Growth and morphology of *E. coli* during tetracycline and erythromycin treatment. Cell growth in bulk culture was measured using a plate reader (A, E). Cell morphology was measured by growing cells on LB agarose pad with 1 μM SYTOX Green and (B-D) 20 μg/mL tetracycline or (F-H) 800 μg/mL erythromycin. A total of 109 and 209 cells for tetracycline and erythromycin, respectively, were used for cell length and mean width measurements, and cell volume was estimated.

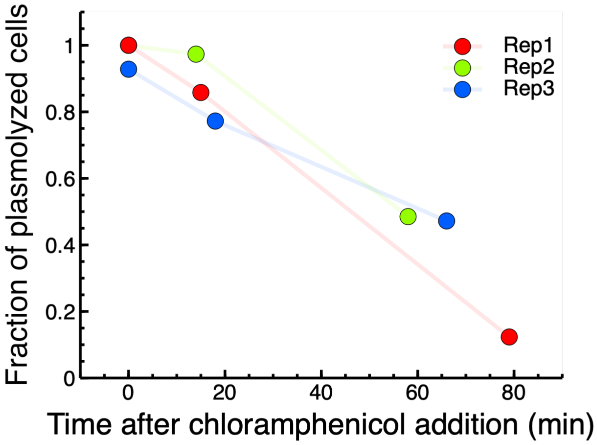


Extended Data Fig 2. Plasmolysis of chloramphenicol treated cells. Chloramphenicol treated cells were incubated on LB agarose pad with 200 mM additional NaCl and imaged using a microscope. 100 – 300 cells per replicate were used for measurement, with total cell counts of 1601.

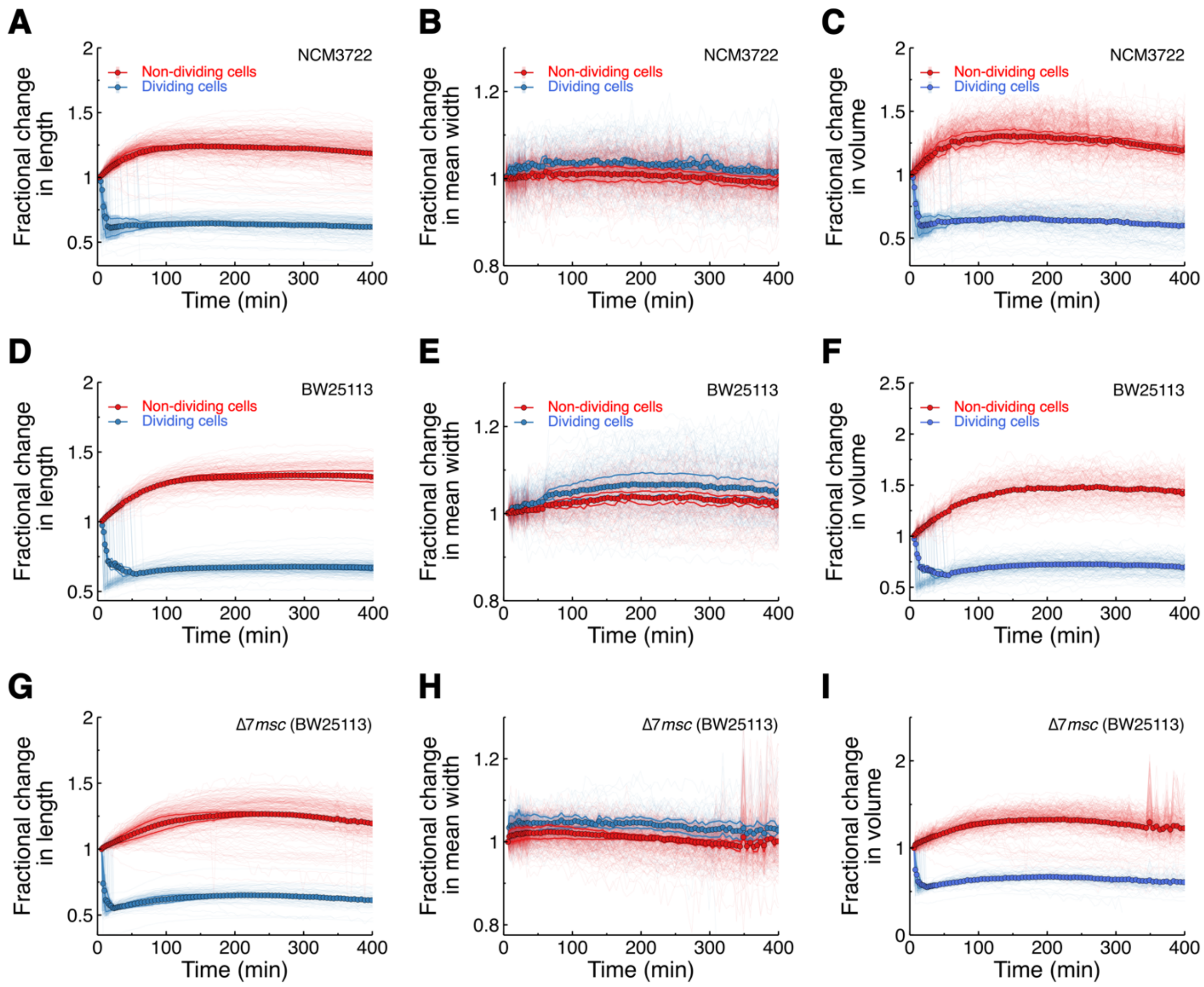


Extended Data Fig 3. Morphology of nutrient-starved cells during chloramphenicol treatment. Log phase cells were incubated on agarose pad prepared in 1.16% NaCl containing 60 μg/mL chloramphenicol and 1 μM SYTOX Green and imaged using a microscope. Cells were separated into dividing and non-dividing populations. Fractional changes in length, mean width, and volume are shown for (A-C) NCM3722, (D-F) BW25113, and (G-I) Δ7*msc*. Two biological replicates were analyzed with total cell counts of 570, 331, and 332, for NCM3722, BW25113, and Δ7*msc*, respectively. Thin lines represent individual cells, bold lines represent the mean of each biological replicate, and markers represent the mean of the two biological replicates.

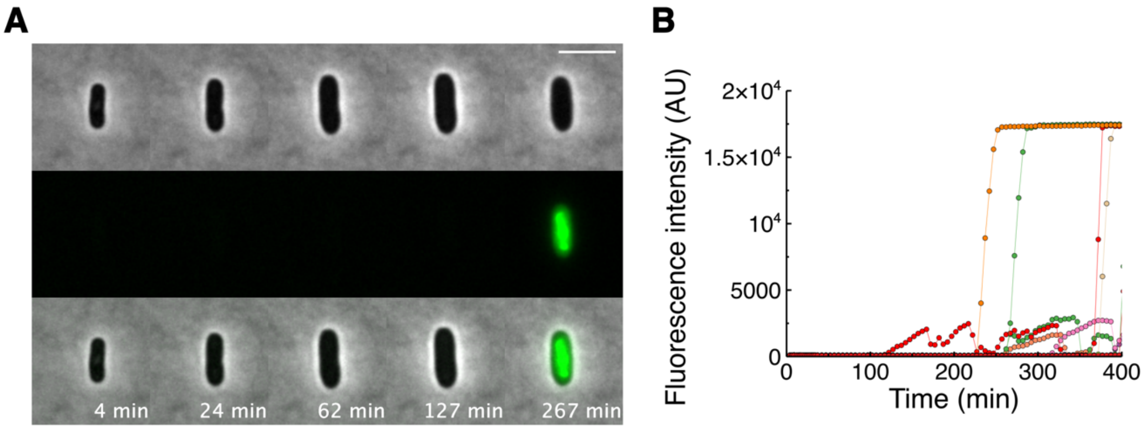


Extended Data Fig 4. Changes in morphology and membrane permeability of a dying cell during chloramphenicol treatment. (A) Representative phase-contrast and SYTOX Green fluorescence images of a cell during chloramphenicol treatment. Cell death was identified by uptake of the membrane-impermeable dye SYTOX Green. Scale bar 5 μm. (B) SYTOX Green fluorescence intensity over time for individual cells. A subset of cells from one biological replicate is shown for clarity.

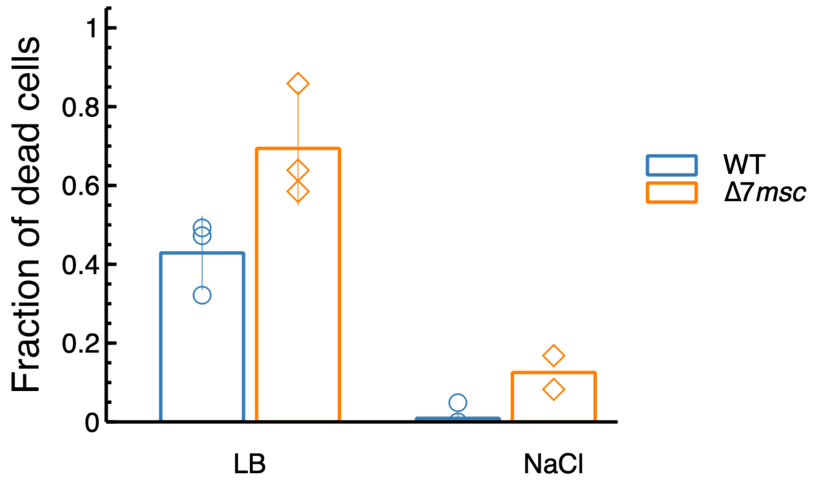


Extended Data Fig 5. Death of nutrient-starved wild type and mechanosensitive channel mutant (Δ7*msc*) cells during chloramphenicol treatment. Log phase cells were incubated on agarose pad prepared in 1.16% NaCl containing 60 μg/mL of chloramphenicol and 1 μM SYTOX Green and imaged by microscopy. A total of 335 wild type and 331 mutant cells from two biological replicates were analyzed for nutrient-starved condition (NaCl) and 630 wild type and 473 mutant cells from three biological replicates for LB. The fraction of dead cells was measured at 300 min of chloramphenicol treatment. LB data are the same as shown in Fig 3C.

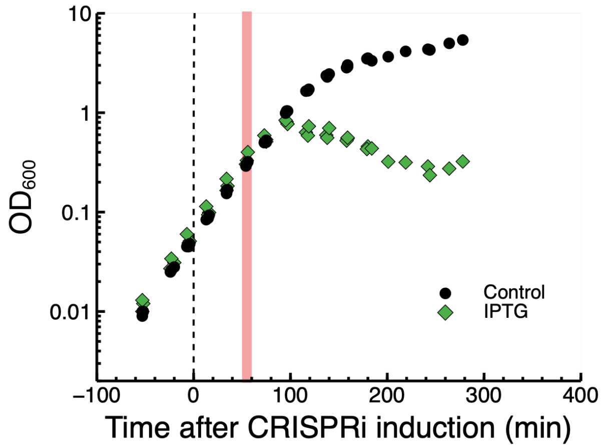


Extended Data Fig 6. Growth of PBP1b-depleting cells in LB. ΔPBP1a strain with chromosomally encoded IPTG-inducible CRISPRi was grown to early log phase ($OD_{600}$ ~0.05, dotted line) and 100 μM of IPTG was added to induce gRNA expression. The red line indicates the time at which chloramphenicol was added in the parallel experiment shown in Fig. 4D.

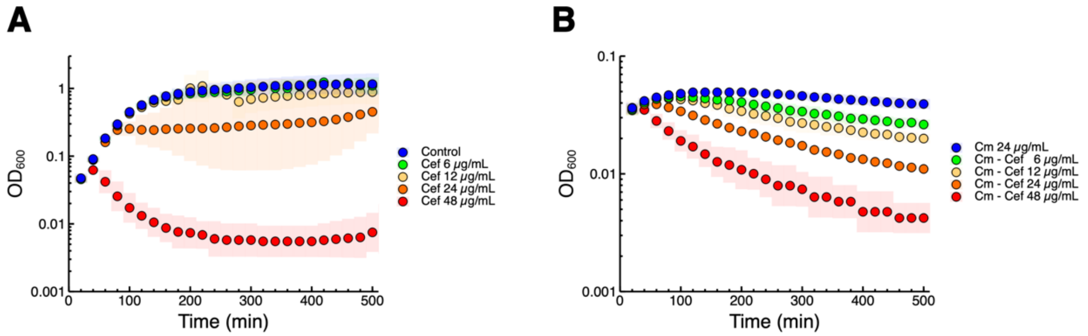


Extended Data Fig 7. Growth of wild type NCM3722 in LB with cefsulodin and chloramphenicol. Exponential phase cells were exposed to single antibiotics or antibiotic combinations on a 96-well plate and $OD_{600}$ was measured using a plate reader every 10 min. Geometric means and standard deviations from three biological replicates are plotted.

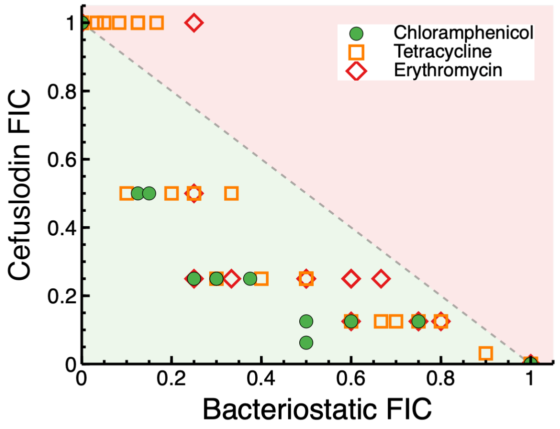


Extended Data Fig 8. Fractional inhibitory concentration (FIC) of bacteriostatic drugs combined with cefsulodin. Exponentially growing cells were plated with combinations of drugs on 96-well plates. Combinations with a summed FIC <1 (green) indicate positive interactions, whereas those with a summed FIC >1 (red) indicate negative interactions.
